# Treatment with carnosic acid attenuates the secretion of pro-inflammatory mast cell mediators following IL-33 activation

**DOI:** 10.1101/2024.09.29.615684

**Authors:** Robert W.E. Crozier, Jordan T. Masi, Adam J. MacNeil

## Abstract

IL-33 is an alarmin cytokine, released upon cellular damage, that has gained significant attention as a regulator of inflammation in several pathologies, including allergy. Once released, IL-33 binds to and activates its cognate receptor ST2, leading to the activation of the classical inflammatory Myddosome signalosome in an array of cells, including mast cells. Our group has recently identified the polyphenol carnosic acid (CA), as a potent regulator of mast cell activation in the context of allergic inflammation. Due to the key role IL-33 plays in the enhancement of allergies and other mast cell-associated diseases, we sought to determine the inhibitory potential of CA in a model of IL-33-mediated mast cell activation. Bone marrow-derived mast cells were stimulated with IL-33 under potentiation of SCF and treated with CA in the presence or absence of an allergen co-stimulation. Here, it was determined that treatment with CA led to a reduction in early ROS production, which translated into a significant impairment in the release of pro-inflammatory cytokines IL-6, IL-13, TNF and chemokines CCL1, CCL2 and CCL3. Surprisingly however, it was determined that CA treatment increased signaling through Akt and NFκB protein phosphorylation, leading to increased gene expression of *IL6, IL13,* and *CCL3* as well as increased intracellular concentrations of IL-6 and CCL3. Taken together, our data suggests treatment with CA impairs the release mechanisms of pro-inflammatory mediators following IL-33 activation, warranting further investigation into the versatile biological activity of CA toward advance our understanding of CA as a potential anti-inflammatory therapeutic.

## Introduction

Interlukin-33 (IL-33) is an alarmin belonging to the IL-1 superfamily of cytokines, which includes IL-1α, IL-β, and IL-18^1^. IL-33 is constitutively sequestered in the nucleus of structural cells exposed to exogenous stimuli such as fibroblasts, endothelial, and epithelial cells, allowing for rapid release following tissue damage and subsequent cell death. IL-33 has demonstrated dual functionality, acting as a nuclear transcription factor by binding to heterochromatin, or functioning as a traditional cytokine following secretion^2^. Initially identified as a potent inducer of Th2 immunity, studies have demonstrated that IL-33 is a highly versatile cytokine capable of inducing eosinophilia and activating mast cells, basophils, CD4+ and CD8+ T cells, innate lymphoid cells, and even monocytes/macrophages^3^. Due to its extensive role in activating a plethora of immune cells, IL-33 finds itself at the center of several inflammatory pathologies, including rheumatoid arthritis, cancer, and cardiovascular disease, and can potentiate the severity of allergic inflammatory disorders, such as asthma and atopic dermatitis^4^.

Once passively released in its pro-peptide form, IL-33 is proteolytically cleaved by local proteases to liberate the chromatin-binding region from the active cytokine domain^4,5^. In its active form, IL-33 binds to its cognate cell surface receptor ST2, where it mediates tissue specific responses such as immune cell recruitment, increased mucus production and the release of inflammatory mediators from local immune cells^6–8^. IL-33-induced signaling initiates ST2 heterodimer formation with the IL-1 receptor accessory protein (IL-1RAcP), which as a complex, lack’s intrinsic enzymatic activity^9^. The ST2 heterodimer complex provides a binding site on its toll-interleukin-1 receptor domain for MyD88, which undergoes oligomerization, and recruits downstream IRAK and TRAF proteins comprising the classical Myddosome signaling complex^8,10^. The upstream phosphorylation cascade initiated through the autophosphorylation of IRAK4, eventually results in the downstream activation of mitogen activated protein kinases (MAPKs), and several transcription factors such as NFκB, all of which are responsible for the production and eventual release of several key inflammatory mediators associated with IL-33-induced inflammation^8^.

Mast cells are densely granulated immune sentinels located within vascularized tissues interfacing the external environment, including the skin, lungs, and gastrointestinal tract^11^. Due to their large receptor repertoire, mast cells have the capability to produce and secrete bioactive mediators in response to various extrinsic stimuli^12^. Accordingly, mast cells initiate and orchestrate innate and adaptive immune responses, thereby contributing to several pathophysiological processes^13^. Mast cells are most notably recognized for their central role in allergic inflammation, as they constitutively express FcεRI, a high-affinity IgE receptor.

Allergen-bound IgE cross-linking of FcεRI triggers mast cell activation, stimulating the release of preformed and newly synthesized inflammatory mediators^14^. Mast cells also express high levels of the cognate IL-33 receptor ST2^15^. *In vitro* studies have shown that IL-33 exacerbates allergen-induced activation of both mouse^16^ and human^14^ mast cells by increasing the production and release of several inflammatory cytokines^2^. Moreover, IL-33 can potently induce mast cell activation and cytokine production even in the absence of IgE-FcεRI stimulation, highlighting its role as a key mediator of mast cell function^16,17^.

IL-33-induced mast cell activation has been implicated the pathogenesis of allergic and non-allergic inflammation, including atopic dermatitis, anaphylaxis, asthma, inflammatory bowel disease, cancers, and several other inflammatory disorders^2,18^. Given that IL-33 expression and protein levels are upregulated in these conditions, modulating the IL-33/ST2 axis is an attractive strategy for therapeutic intervention^12^. Recently, natural compounds derived from plants have received considerable attention due to their low toxicity and extensive pharmacological properties^19^. As one of the most abundant secondary metabolites found in fruits and vegetables, polyphenols are well known for their anti-oxidant, anti-tumor, anti-bacterial, and anti-inflammatory properties, and are thus an important source of novel therapeutics^20^. We recently demonstrated that carnosic acid (CA), an abundant polyphenolic constituent found in *Rosmarinus officinalis* L., inhibits IgE-mediated mast cell activation by modulating downstream signaling pathways, resulting in impaired degranulation and reduced pro-inflammatory mediator production and release^21^. In the present study, we found that CA potently attenuates IL-33-mediated mast cell secretion of pro-inflammatory mediators in the presence and absence of FcεRI co-stimulation, suggesting the potential use of CA as a therapeutic for treating allergic and other IL-33-mediated inflammatory diseases.

## Materials and Methods

### Mice

Female C56BL/6 mice (Charles River Laboratories) were housed and maintained under standard chow diet and living conditions. All protocols were approved by the Animal Care Committee at Brock University in accordance with The Canadian Council on Animal Care guidelines. AUP: 20-07-02.

### Bone marrow-derived mast cells

Bone marrow was isolated from the femur and tibia of C57BL/6 mice and hematopoietic stem and progenitor cells were flushed. Freshly isolated stem and progenitor cells were differentiated in the presence of IL-3 (10% WEHI-3B supernatant)-conditioned media comprised of RPMI-1640 (Gibco, 11875-093), 10% Serum Plus II (Sigma, 14009C), 1% pen/strep (Sigma, P0781), 200 nM PGE2 (Sigma, P5640) and 50 nM 2-mercaptoethanol (Sigma, M7522). BMMC complete media was changed every 3 to 4 days and cultures were maintained at 37 °C with 5% CO2. Following the 5-week differentiation and growth period, mast cell maturity was assessed *via* flow cytometry and determined by the percentage of FcεRI^+^/c-kit^+^ cells in culture. Following the differentiation period, BMMC cultures were maintained in the BMMC complete media described above.

### Mast cell sensitization, stimulation and polyphenol treatment

Regardless of the stimulation conditions, mast cells were sensitized overnight with allergen specific IgE antibodies as previously described^21^. The following day, BMMCs were washed twice with sterile RPMI-1640 to remove any unbound antibodies. Cells were then stimulated with either 50 ng/ml mIL-33 (Biolegend, 580506) under potentiation of 100 ng/ml SCF (PeproTech Inc., 250-03) or mast cells were stimulated with 100 ng/ml TNP-BSA (Biosearch Technologies, T-5050-10), 100 ng/ml SCF and 50 ng/ml mIL-33. BMMCs were simultaneously treated with 50 μM CA (Sigma, C0609) or equal volume of DMSO as a vehicle control.

### Flow Cytometry

For cell surface receptor analysis, BMMCs were centrifuged at 340 ×*g* for 10 mins and washed with 6 ml immunofluorescence buffer (IF) (PBS with 0.2% NaN3 and 1% BSA). Cells were resuspended at 0.5 x10^6^/ml and 100 μl of cells was stained per condition on ice for 1 hour with fluorochrome-conjugated antibodies; PE-conjugated anti-mouse FcεRIα (#1271535), Armenian hamster isotype control (#2604535), FITC-conjugated anti-mouse CD117 (c-kit) (#1129025) and rat IgG2b (#2603025) isotype control from Sony Biotechnology and APC-conjugated anti-IL-33Ra (ST2) (#146605) and Rat IgG1, κ isotype control (#400411) from BioLegend. Stained BMMCs were washed twice with 600 μL IF buffer and fixed in 400 μL of 0.37% formalin. For ROS analysis, BMMCs were stained with 5 μM 2’, 7’-DCFH-DA (Sigma, D6883) for 30 mins at 37 °C. BMMCs were then treated with CA (Sigma, C0609) and simultaneously stimulated with 50 ng/ml mIL-33 and 100 ng/ml SCF for 15 mins, placed on ice immediately following incubation and washed twice with HBSS (Sigma, 55037C). All data were acquired on a Sony SH800S cell sorter (Sony Biotechnology).

### β-hexosaminidase release assay

BMMCs were sensitized overnight and washed above as previously described with cold Hanks’ balanced salt solution (HBSS) supplemented with 0.1% BSA used in placed of RPMI-1640. Cells were resuspended at a density of 4 x10^6^ cells/ml and aliquoted in duplicate in a 96-well tissue culture plate and stimulated with 50 ng/ml IL-33 and 100 ng/ml SCF, 100 ng/ml TNP-BSA and 100 ng/ml SCF or 50 ng/ml IL-33, 100 ng/ml TNP-BSA and 100 ng/ml SCF with or without 50 μM CA treatment for 30 mins.

### Enzyme-linked immunosorbent assay

For measurement of secreted cytokine and chemokines following IL-33 stimulation, BMMCs were stimulated with 50 ng/ml mIL-33 and 100 ng/ml SCF at a density of 2 x10^6^ cells/ml for 1, 6 and 24 hours. At each timepoint, cells were centrifuged at 1152*g* for 10 min at 4 °C to collect the supernatant, which was subsequently stored at -80 °C until ready to analyze.

For measurement of intracellular cytokine and chemokine concentrations, BMMCs were stimulated with 50 ng/ml mIL-33 and 100 ng/ml SCF for 6 and 24 hours. At each timepoint, samples were centrifuged at 1152*g* for 10 min at 4 °C to pellet the cells. The supernatant was removed and stored for future analysis, while the pellet was washed with PBS. Following the wash, pellets were lysed in 60 μL of extraction buffer (100 mM Tris, 150 mM NaCl, 1 mM EGTA, 1 mM EDTA, 1% Triton X-100, 1 mM PMSF and 80 uL of PPI cocktail Sigma, P8340) and total sample protein was determined with a Bradford assay. ELISA assays were performed using DuoSet kits (R&D IL-6 DY406, IL-13 DY413, TNF DY410, CCL1 DY845, CCL2 DY479 and CCL3 DY450) according to the manufacturer’s instructions. Assays were performed on Nunc MaxiSorp flat bottom 96-well plates (Thermo-Fisher, 44-2404-21) and developed with the addition of 100 μL of BD OptEIA TMB Substrate (BD Biosciences, 555214). Optical density was measured using a spectrophotometer (BIO-TEK) at a 450 nm wavelength. Following development of each ELISA plate, calculated cytokine/chemokine concentrations were normalized to the total cellular protein concentration of the corresponding sample that was determined with a Bradford assay.

*RNA isolation and qPCR* – RNA was isolated from 3 x10^6^ BMMCs using the RNeasy Plus kit (Qiagen, 74136) according to the manufacturer’s instructions. Isolated RNA was quantified on a NanoVue spectrophotometer (GE) and cDNA reactions were prepared using 500 ng of RNA with double-primer RNA to cDNA premix (TakaraBio, 63549) and diluted to a 1:20 working stock in molecular grade H2O (Sigma, 693520). qPCR assays were performed on an ABI StepOnePlus Real-Time PCR instrument (Applied Biosystems, 4376599) with KAPA SYBR fast master mix (Sigma, KM4113) and amplification efficiency-optimized primers (Integrated DNA Technologies). Primers used included *IL6 For 5’ AGACAAAGCCAGAGTCCTTCAGAGA 3’ Rev 5’ TGGTCTTGGTCCTTAGCCACTCC 3’, IL13 For 5’ TGCTTGCCTTGGTGGTCTCG 3’ Rev 5’ TCCATACCATGCTGCCGTTGC 3’, TNF For 5’ TGAACTTCGGGGTGATCGGTCC 3’ Rev 5’ TCCAGCTGCTCCTCCACTTGGT 3’, CCL1 For 5’ CCAGACATTCGGCGGTTGCT 3’ Rev 5 CAAGCAGCAGCTATTGGAGACCGT 3’, CCL2 For 5’ CCTCCACCACCATGCAGGTCC 3’ Rev 5’ CAGCAGGTGAGTGGGGCGTTA 3’, CCL3 For 5’ GGAGCTGACACCCCGACTGC 3’ Rev 5’ GGGTTCCTCGCTGCCTCCAA 3’ and HPRT For 5’ CTTGCTGGTGAAAAGGACCTCTCG 3’, Rev 5’ CGCTCATCTTAGGCTTTGTATTTGG 3’*.

Threshold cycle (CT) values were recorded and analyzed using the ΔΔCt method to determine the fold change of expression of a gene of interest relative to the reference gene hypoxanthine-guanine phosphoribosyltransferase (*hprt*).

#### Protein extraction and western blot

BMMCs (5-10 x10^6^) were lysed with radioimmunoprecipitation buffer (Sigma, R0278) supplemented with 10 μg/ml leupeptin (Bio Basic Canada Inc., AD0153), 10 μg/ml pepstatin (Bio Basic Canada Inc., PPDJ694), 10 μg/ml aprotinin (Bio Basic Canada Inc., AD0153), 0.2 mM iodoacetamide (Bio Basic Canada Inc., IB0539), 1 mM NaF (Sigma, S-7929), 1 mM Na3VO4 (Sigma, S-6508), 0.5 mM PMSF (Bio Basic Canada Inc., PB0425) and PIC (Sigma, P8340). Purified lysates were loaded onto 10% Tris-glycine acrylamide gels (BioRad, 1610156) and electrophoresed for ∼40 mins at 200 V. Separated proteins were transferred onto 0.2 μm PVDF membranes *via* transblot turbo (BioRad, 1704150) and subsequently blocked with 5% non-fat milk or BSA and probed with primary antibodies from Cell Signaling Technologies against phospho-proteins p-Akt (Ser473, 4060) p-ERK1/2 (Thr202/Tyr204, 9101), p-IRAK4 (Thr435/Ser346, 11927), p-IκBα (Ser32/36, 9246), p-JNK (Thr183/Tyr185, 9211), p-p38 (Thr180/Tyr182, 9251) and total-proteins Akt (4691), ERK1/2 (4695), JNK (9252), IRAK4 (4363), Myd88 (4283), p38 (8690) and Vinculin (Millipore-Sigma, 05-386). Goat anti-rabbit (7074) and horse anti-mouse (7076) HRP-conjugated secondary antibodies were purchased from Cell Signaling Technologies. Clarity max ECL substrate (BioRad, 170-5062) was applied to membranes and images were visualized on a BioRad ChemiDoc imager (BioRad, 12003153) and band intensity was quantified using the corresponding BioRad Image Lab software (https://www.bio-rad.com/en-ca/product/image-lab-software?ID-KRE6P5E8Z).

#### Statistical Analysis

Results were expressed as mean ± SEM and were analyzed using a two-tailed paired *t* test or two-way ANOVA with a Dunnet’s *post-hoc* test where appropriate.

Statistical analyses were conducted using GraphPad Prism (v.10.0.2) and results were reported as statistically significant if *p* <0.05.

## Results

### Carnosic acid treatment exhibits antioxidant properties following IL-33 activation

Our previous work looking at the effects of CA in a mouse mast cell model of allergic inflammation concluded that treatment of BMMCs with 50 μM CA resulted in a significant downregulation of IgE-FcεRI complexes in addition to c-kit receptor levels on the cell surface. Here, we sought to determine if the effects of CA on downregulating cell surface receptor levels was specific to inherent biochemistry and/or cycling kinetics of receptors involved in allergy activation, or if it is a widespread effect observed at the cell surface following treatment with CA. To investigate this angle, BMMCs were treated with 50 μM CA and stimulated with IL-33 and SCF for 30 mins prior to staining with a fluorescent antibody specific to IL-33Rα (ST2).

Although a slight inhibitory trend exists, it was determined that treatment with CA does not alter levels of surface ST2 (Fig. 1A-B). These results were consistent with mRNA levels for *IL1rl1*, the encoding gene for ST2, which remained unchanged following CA treatment and IL-33 stimulation up to 360 mins (Fig. 1C), further indicating that any downstream inhibitory effects observed are the result of impairments to molecular mechanisms, and not disruption of cell activation at the cell surface level.

**Figure 1.**
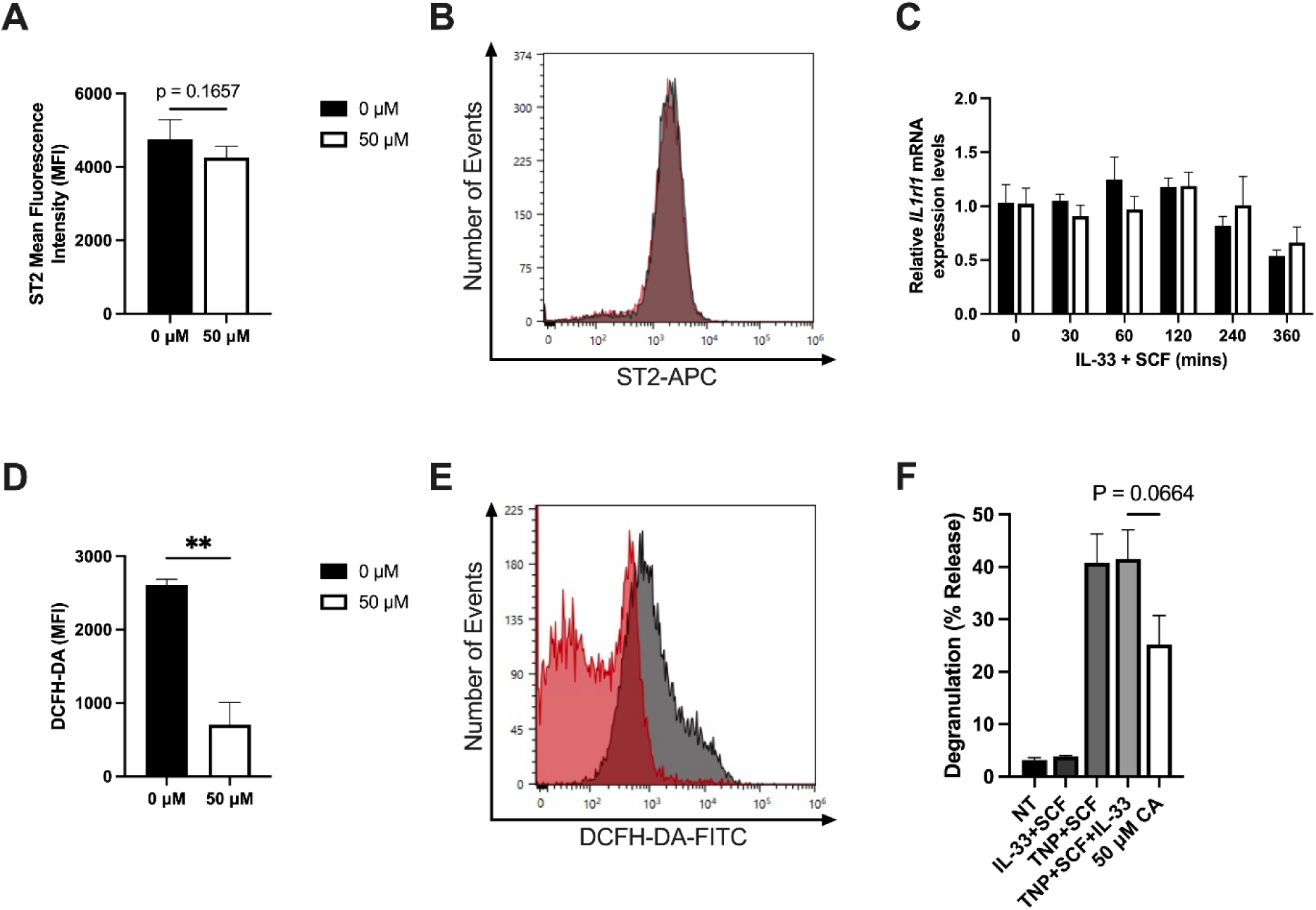
Treatment with CA impairs IL-33-mediated ROS production. (A-B) BMMCs were treated with 50 μM CA and simultaneously stimulated with 50 ng/ml IL-33 and 100 ng/ml SCF for 30 mins. Following stimulation, BMMCs were stained with APC-conjugated anti-mouse ST2 antibodies to look at surface level receptor presence. Data expressed as mean ±SEM of 3 independent primary BMMC cultures, and a two-tailed paired t-test was used to determine any differences between 0 and 50 μM CA. **(C)** CA treated BMMCs were stimulated with IL-33 for 0, 30, 60, 120, 240 and 360 mins. Isolated mRNA was used to quantify the relative gene expression levels of *IL1rl.* Data expressed as mean ±SEM of 3 independent primary BMMC cultures. A two-way ANOVA and Dunnett’s multiple comparison were used to determine differences between 0 and 50 μM CA. **(D-E)** BMMCs were stained with 5 μM DCFH-DA for 30 mins, stimulated with IL-33 and SCF and treated with 50 μM CA for 15 mins to measure ROS production. Data expressed as mean ±SEM of 3 independent primary BMMC cultures. A two-tailed paired t-test was used to determine any differences between 0 and 50 μM CA. ** p<0.01. **(F)** BMMCs were stimulated in the presence of IL-33+SCF, TNP+SCF, TNP+SCF+IL-33 and TNP+SCF+IL-33 following treatment with CA prior to measure the release of β-hexosaminidase following stimulation. Data expressed as mean ±SEM of 3 independent primary BMMC cultures. A one-way ANOVA was used to determine any differences between 0 and 50 μM CA.

We next wanted to determine the antioxidant effects of CA treatment following IL-33 induced mast cell activation. BMMCs were pre-treated with 5 μM 2’, 7’-DCFH-DA for 30 mins. Following incubation, BMMCs were treated with 50 μM of CA and simultaneously stimulated with 50 ng/ml mIL-33 and 100 ng/ml SCF for 15 mins, and subsequently analyzed via flow cytometry. As expected, it was observed that CA treatment led to a significant reduction of IL-33-induced ROS production, which was observed by a significant reduction of DCF-associated MFI when compared to the control (Fig. 1D-E). Although unable to induce the mobilization and the subsequent degranulation response when stimulated alone, the presence of IL-33 in the context of allergy has been shown to potentiate degranulation when mast cells are activated with IL-33 alongside allergen and SCF. As a result, we examined if stimulation with 50 ng/ml IL-33, 100 ng/ml TNP-BSA and 100 ng/ml SCF would lead to increased degranulation and if treatment with 50 μM CA would still be an effective inhibitor of the early phase degranulation response. Here, we observed that IL-33 stimulation alongside TNP-BSA and SCF did not increase the degranulation response compared to TNP-BSA + SCF alone, contrary to previous reports.

Additionally, we found that treatment with 50 μM CA was able to successfully reduce the release of mast cell granules in the presence of all 3 stimuli (Fig. 1F). Taken together, these results show that CA treatment impairs early molecular events following IL-33-mediated mast cell activation (i.e., ROS production), independent of any modulation to functional IL-33R located on the cell surface.

### Carnosic acid impairs IL-33-induced and IL-33 potentiated cytokine and chemokine release

Activation of immune cells through the Myddosome signaling network *via* various stimuli (TLRs, cytokine receptors) has been well characterized as a potent inducer of cytokine and chemokine release^10^. IL-33-mediated mast cell activation during various contexts (allergy, local inflammation etc.) shares activation of the Myddosome signalosome, and has been found to drive the production and release of key mediators such as IL-6, IL-13, TNF, CCL2 ^22,23^. Here, BMMCs were stimulated with 50 ng/ml IL-33 and 100 ng/ml of SCF for 6 and 24 hours to determine the effects of CA treatment on IL-33-induced mediator release *via* analyte specific ELISA experiments. Following activation, it was determined that 50 μM CA significantly impaired the release of critical cytokines IL-6 at 24 hr (Fig. 2A), IL-13 at 6 and 24 hr (Fig. 2B) and TNF at 24 hr (Fig. 2C). These results were accompanied by a significant reduction in the release of key chemokines CCL1 (Fig. 2D), CCL2 (Fig. 2E) and CCL3 (Fig. 2F) following 24 hrs.

**Figure 2.**
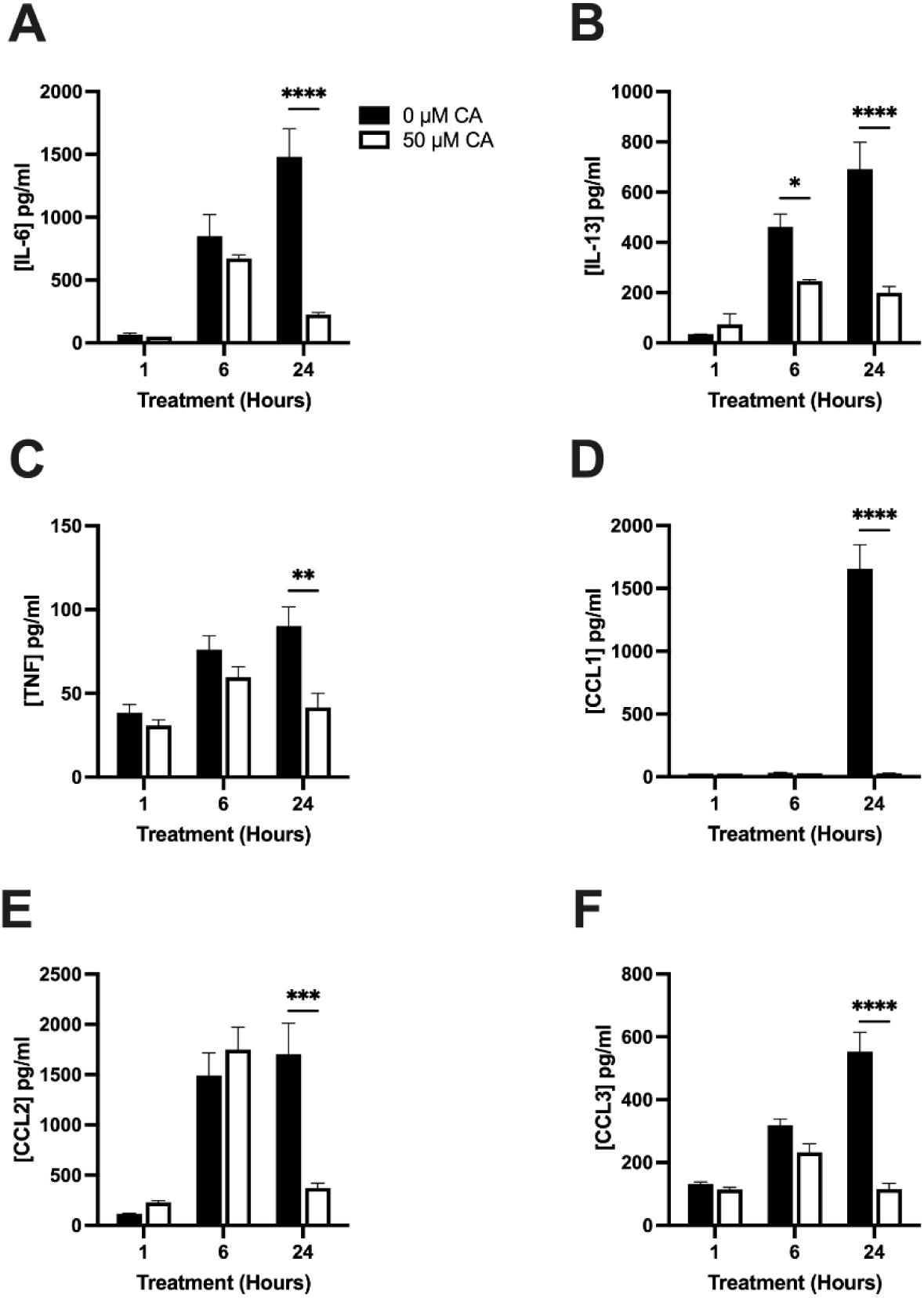
Carnosic acid impairs IL-33-mediated mast cell cytokine and chemokine release. BMMCs (2x 10^6^) were treated with CA and simultaneously stimulated with 50 ng/ml IL-33 under potentiation of 100 ng/ml SCF for 1, 6, and 24 hours. At each timepoint, supernatant was collected and analyzed via ELISA to measure release concentrations of **(A)** IL-6, **(B)** IL-13, **(C)** TNF, **(D)** CCL1, **(E)** CCL2 and **(F)** CCL3. Data expressed as mean ±SEM of 3 independent primary BMMC cultures. A two-way ANOVA and Dunnett’s multiple comparison were used to determine differences in cytokine and chemokine release throughout the stimulation time course following CA treatment. * p<0.05, ** p<0.01, *** p<0.001, **** p<0.0001 relative to BMMCs not treated with CA.

In addition to being able to independently activate immune cells in various pathological contexts, IL-33 has previously been found to synergistically enhance cytokine release in allergen/antigen stimulated mast cells^23^. As a result, we wanted to determine if CA was able to combat cytokine/chemokine release following co-stimulation with allergen (TNP-BSA), SCF and IL-33. Here it was observed that stimulation with TNP-BSA + SCF + IL-33 resulted in a drastic increase of IL-6, IL-13, TNF, CCL1, CCL2 and CCL3 compared to IL-33 + mSCF and TNP-BSA + SCF, indicating that IL-33 does amplify the inflammatory response at sites of allergen exposure. Following CA treatment, a similar inhibitory effect was observed which resulted in the impairment of IL-6 (Fig. 3A) and IL-13 (Fig 3B) release at 6 and 24 hr, TNF at 6 hr (Fig. 3C), CLL1 at 6 and 24 hr (Fig. 3D) in addition to CCL2 (Fig. 3E) and CCL3 at 24 hr (Fig. 3F).

**Figure 3.**
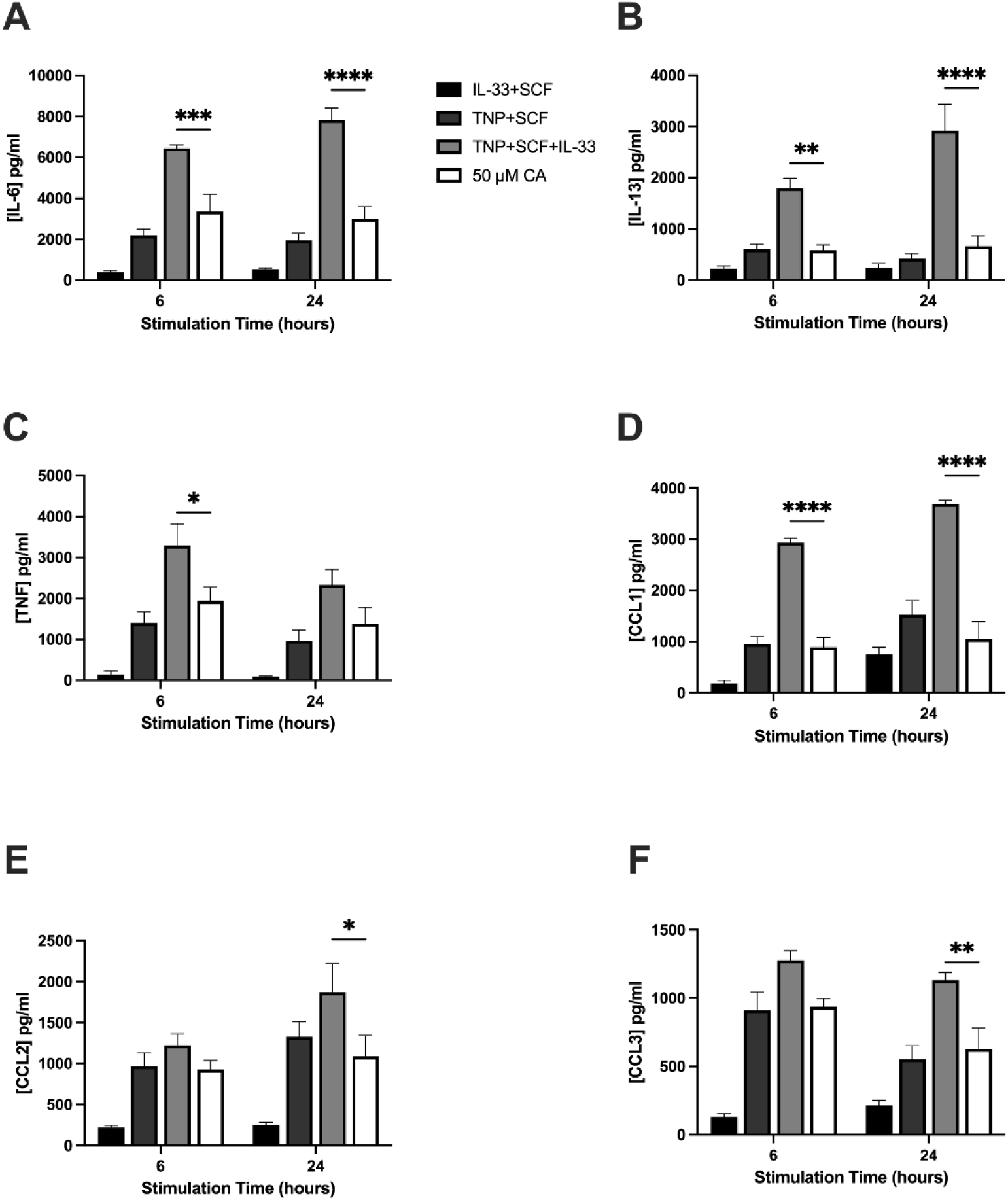
Impairment of IL-33 potentiated cytokine and chemokine release following allergen activation and CA treatment. BMMCs (2x 10^6^) were treated with and without CA and simultaneously stimulated with 50 ng/ml IL-33, 100 ng/ml TNP-BSA and 100 ng/ml SCF for 6, and 24 hours. At each timepoint, supernatant was collected and analyzed via ELISA to measure release concentrations of **(A)** IL-6, **(B)** IL-13, **(C)** TNF, **(D)** CCL1, **(E)** CCL2 and **(F)** CCL3. Data expressed as mean ±SEM of 3 independent primary BMMC cultures. A two-way ANOVA and Dunnett’s multiple comparison were used to determine differences in cytokine and chemokine release throughout the stimulation time course following CA treatment and co-stimulation with IL-33, SCF and TNP-BSA. * p<0.05, ** p<0.01, *** p<0.001, **** p<0.0001 relative to BMMCs not treated with CA.

### Carnosic acid increases gene expression and production of key inflammatory mediators

IL-33-mediated production and release of pro-inflammatory mediators in mast cells is dependent on the activation of several downstream transcription factors such as c-jun, ATF2, NFAT and NFκB^23^. Previously, our group identified that CA-mediated inhibition of inflammatory cytokine and chemokine release following IgE-mast cell activation was dependent on the impairment of upstream gene expression^21^. Here, mast cells were stimulated with IL-33 and SCF for 0, 30, 60, 120, 240, and 360 mins to better understand the underlying mechanism of CA treatment.

Unexpectedly, it was observed that treatment with CA induced increased gene expression of both *IL6* and *IL13* at 240 mins following IL-33 and SCF activation (Fig. 4A-B). Although there was increased of *TNF* gene expression at 240 mins that was non-significant, CA treatment did not influence mRNA levels of *TNF*, *CCL1* or *CCL2* (Fig 4C-E). Similarly, to *IL6* and *IL13* however, treatment with CA was also responsible for a significant increase in *CCL3* gene expression at 240 mins (Fig. 4F).

**Figure 4.**
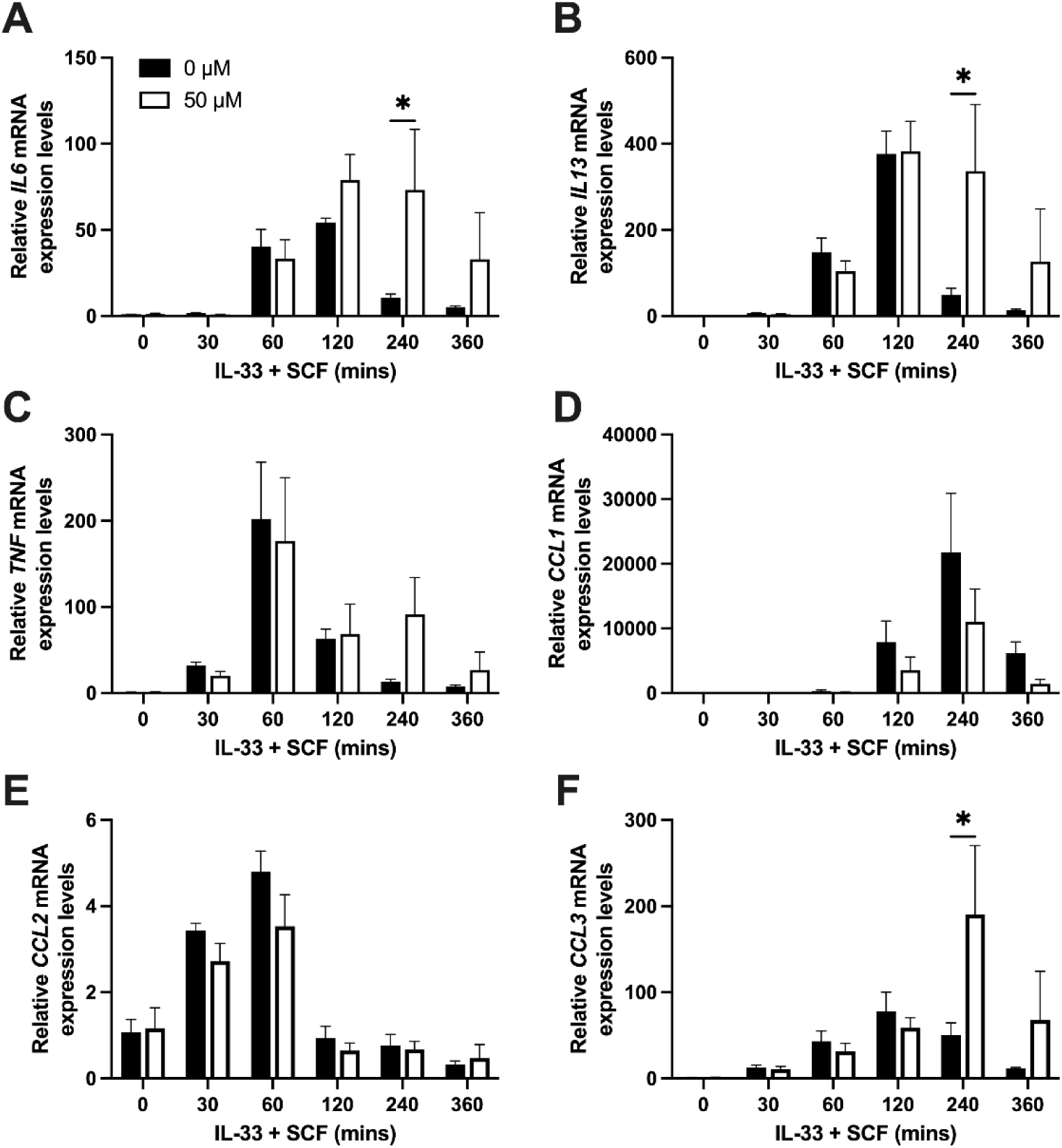
CA treatment increases IL-33-SCF pro-inflammatory gene expression. Isolated mRNA from BMMCs stimulated with IL-33 + SCF for 0, 30, 60, 120, 240 and 300 mins in the presence or absence of CA was analyzed by qPCR to quantify gene expression changes of **(A)** IL6 **(B)** IL13, **(C)** TNF, **(D)** CCL2, **(E)** CCL2, and **(F)** CCL3. Data expressed as mean fold change ±SEM of 3 independent primary BMMC cultures. A two-way ANOVA and Dunnett’s multiple comparison were used to determine differences in cytokine and chemokine gene expression throughout the stimulation time course following CA treatment. * p<0.05 relative to BMMCs not treated with CA (0 μM).

Following these contradictory results with regards to the effect of CA on cytokine and chemokine release, we pursued where exactly CA was having an interruptive role within the protein translation and transportation pathways. To address this concern, BMMCs were stimulated with IL-33 and SCF while simultaneously treated with CA for 6 and 24 hrs to determine intracellular cytokine and chemokine concentrations. Consistent with the gene expression findings in Fig. 4, although not statistically significant, treatment with CA lead to an increase in intracellular IL-6 levels (p=0.07) at 6 hrs, with an increasing trend following 24 hrs (Fig. 5A). Interestingly however, intracellular cytokine levels remained unchanged for IL-13 (Fig. 5B) despite a significant increase of mRNA levels (Fig. 4B). Intracellular protein levels of TNF (Fig. 5C), CCL1 (Fig. 5D) and CCL2 (Fig. 5E) remained unchanged and consistent with their mRNA levels. Expectedly however, intracellular CCL3 levels were significantly increased at 24 hrs (Fig. 5F), corresponding with the significant mRNA levels that were observed at 240 mins and the increasing trend at 360 mins (Fig. 4F). Taken together with the results from Fig. 4, here we show that treatment with CA increases the mechanisms responsible for the production of select proteins, despite inhibiting their ability to be released from IL-33 activated mast cells.

**Figure 5.**
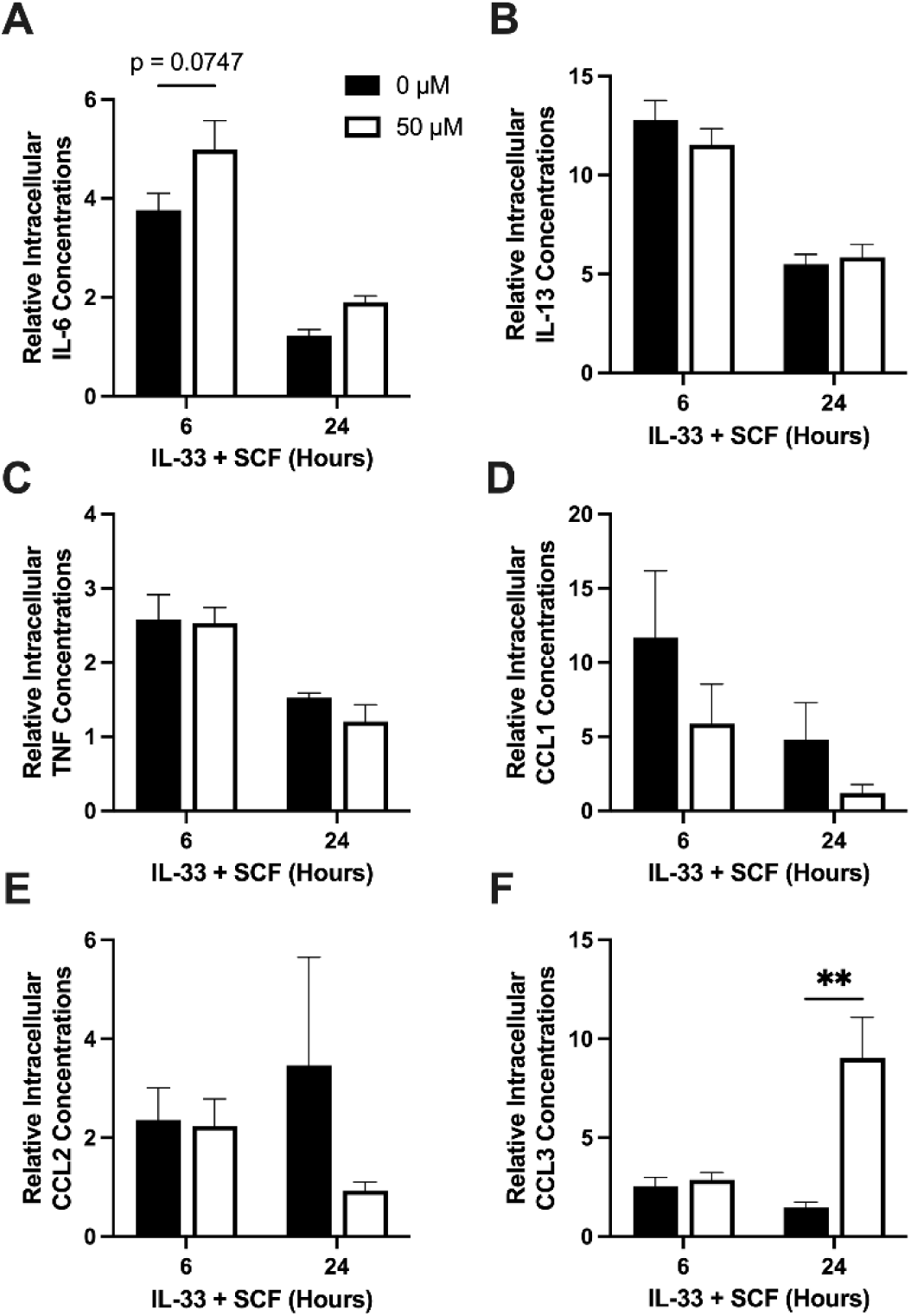
Intracellular mediator concentrations are increased following treatment with CA. IgE-sensitized BMMCs (2x 10^6^) were treated with CA and simultaneously stimulated with 50 ng/ml IL-33, 100 ng/ml TNP-BSA and 100 ng/ml SCF for 6, and 24 hours. At each timepoint, supernatant was decanted, and cell pellets were washed, subsequently lysed and remaining cell debris was cleared. Cell lysates were diluted 1:10 and analyzed via ELISA to measure intracellular concentrations of **(A)** IL-6, **(B)** IL-13, **(C)** TNF, **(D)** CCL1, **(E)** CCL2 and **(F)** CCL3. Data expressed as mean ±SEM of 3 independent primary BMMC cultures. A two-way ANOVA and Dunnett’s multiple comparison were used to determine differences in cytokine and chemokine release throughout the stimulation time course following CA treatment. ** p<0.01 relative to BMMCs not treated with CA.

### Carnosic acid treatment increases key inflammatory pathways Akt and NFκB following IL-33 stimulation

Prior to the production and release of several key inflammatory mediators outlined above, IL-33-ST2 interaction initiates the activation of an expansive signaling network driven through several well characterized nodes such as the Myddosome, MAPK, Akt and NFκB cascades^8^. To dissect the mechanism of action behind the CA inhibitory effect on mediator release but increased gene expression, we investigated protein changes of key players of the Myddosome signaling network, the MAPK, Akt and NFκB pathway following 0-, 5-, 20- and 60-mins IL-33 activation and CA treatment (Fig. 6A). Here it was discovered that directly downstream of the ST2-IL-1RAcP signaling complex, an increasing trend of relative Myd88 protein was observed at 60 mins following treatment with CA (Fig. 6B). Additionally, although an overall CA treatment inhibitory effect on TAK1 phosphorylation (S412) was observed (Fig. 6B), downstream MAPK proteins ERK, JNK and p38 remained unaffected (Fig. 6C-E), apart from an increase in p38 phosphorylation at 0 mins following treatment with 50 μM CA (Fig. 6E). The effect of CA on MAPK signaling in our mast cell model is consistent with what was observed previously when BMMCs were treated with CA and stimulated with allergen^21^. Differences from our previous investigation were observed however when looking at the effects of CA on Akt and upstream regulators of the NFκB pathway (Fig. 7A). Here, we discovered that CA resulted in a statistically significant increase in the overall treatment effect of CA (p=0.020) when comparing phosphorylation of Akt (S473) to our untreated BMMCs, with an increasing trend observed at 20 mins (Fig. 7B). This result was coupled with a significant increase in the phosphorylation status of the NFκB regulator IκBα at 5 mins following CA treatment and IL-33 activation (Fig. 7B).

**Figure 6.**
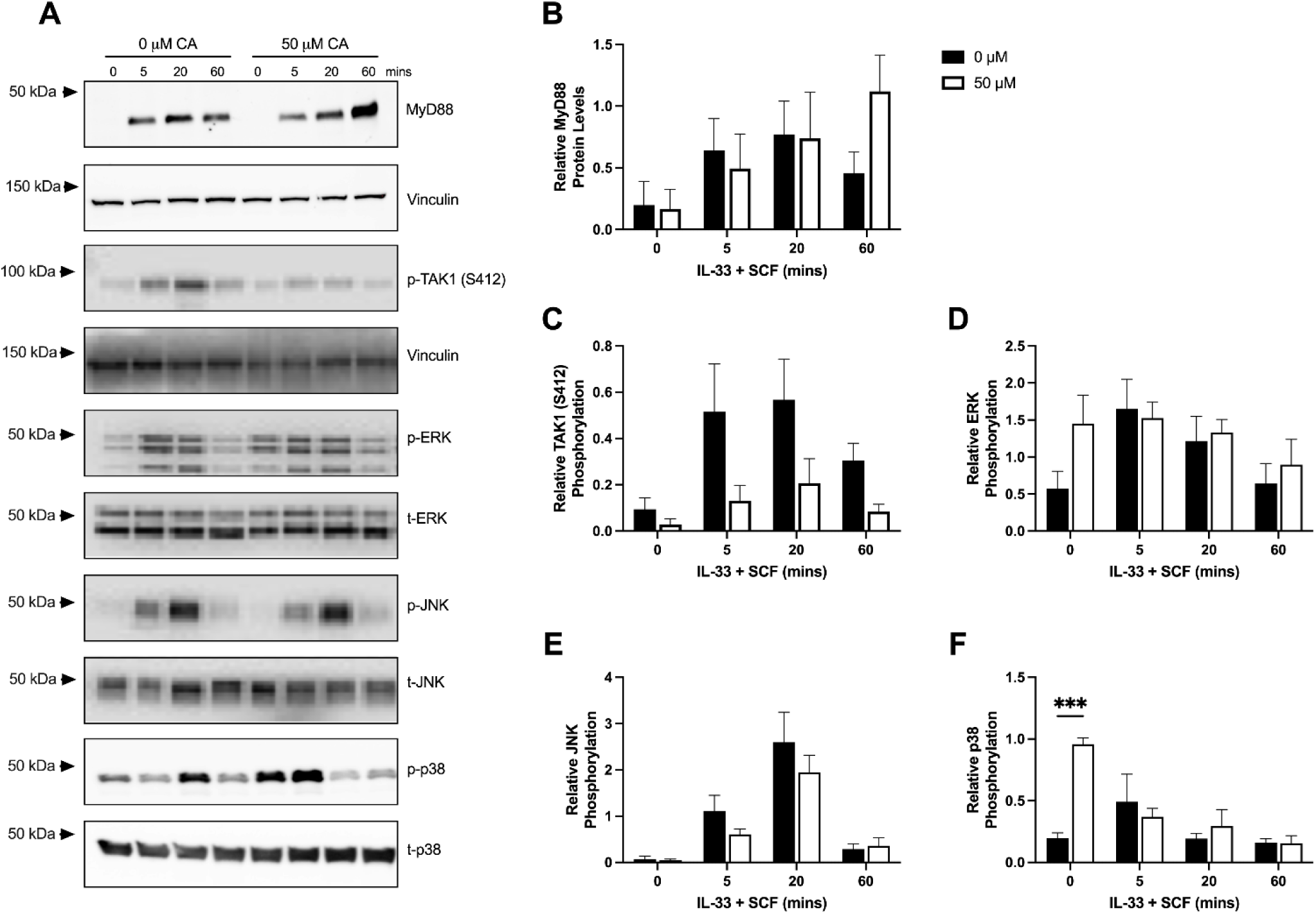
Treatment with CA has differential effects on key signaling nodes. Protein isolates from untreated and CA treated BMMCs stimulated with IL-33 + SCF for 5, 20 and 60 mins were analyzed via western blot. **(A)** Representative blots of n=3 independent primary BMMC cultures showing MyD88, vinculin, p-TAK1, p- and t-ERK, p- and t-JNK and p- and t-p38. Relative protein levels (total protein/Vinculin) were quantified for **(B)** MyD88, while relative phosphorylation (phosphor/total) was quantified for **(C)** TAK1 (S412**), (D)** ERK, **(E)** JNK, and **(F)** p38, in the presence or absence of CA treatment. Data expressed as mean relative phosphorylation ±SEM of 3 independent primary BMMC cultures. A two-way ANOVA and Dunnett’s multiple comparison were used to determine differences in relative phosphorylation levels following allergen stimulation and CA treatment. *** p<0.001 relative to BMMCs not treated with CA.

**Figure 7.**
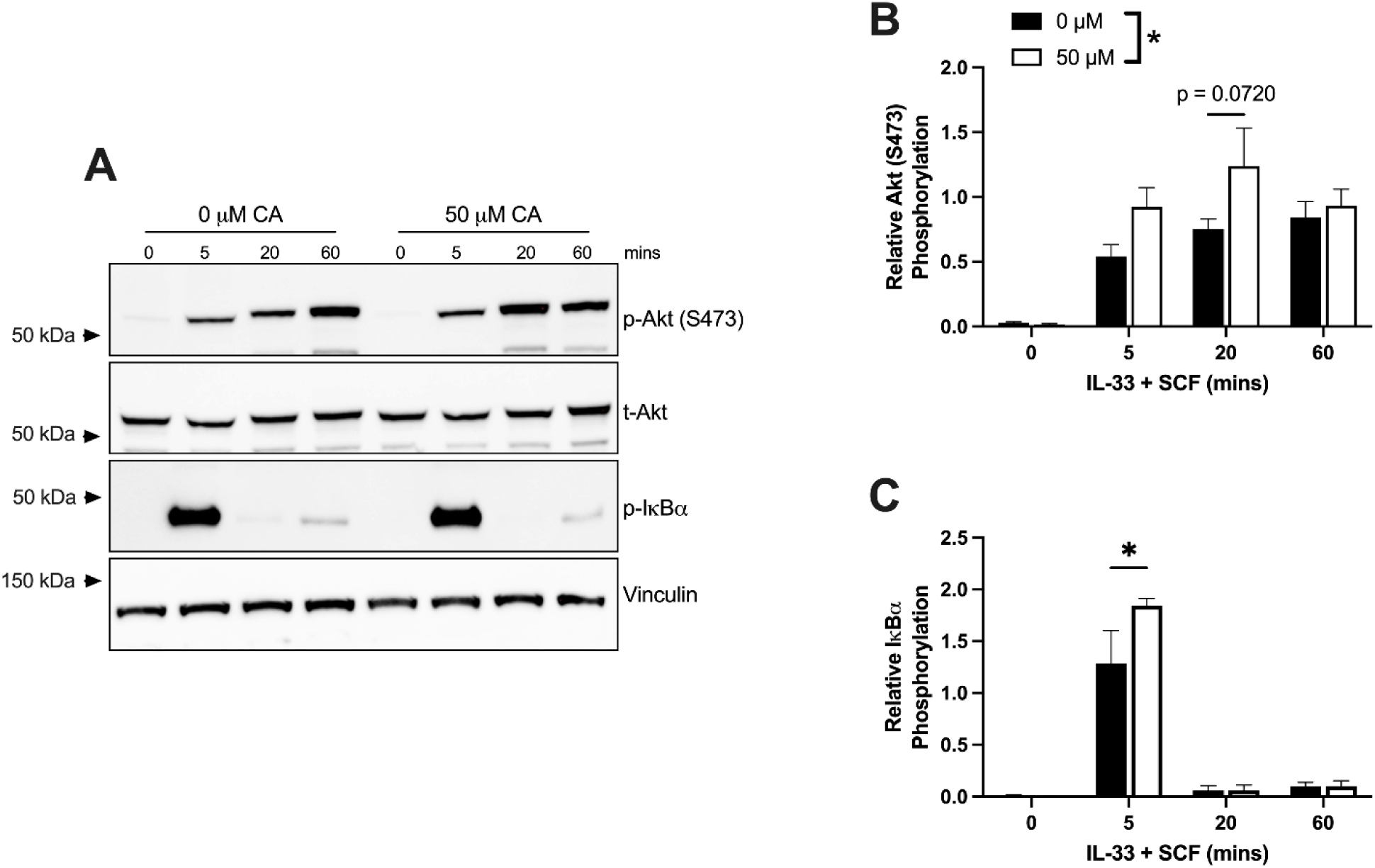
IL-33-induced Akt and NFκB signaling is increased following CA treatment. Protein isolated from untreated and CA treated BMMCs stimulated with IL-33 + SCF for 5, 20 and 60 mins were analyzed via western blot. **(A)** Representative blots of n=3 independent primary BMMC cultures showing p- and t-Akt, p-IκBα and vinculin. Relative protein levels (phospho protein/Vinculin) were quantified for **(B)** Akt (473) and **(C)** IκBα in the presence or absence of CA treatment. Data expressed as mean relative phosphorylation ±SEM of 3 independent primary BMMC cultures. A two-way ANOVA and Dunnett’s multiple comparison were used to determine differences in relative phosphorylation levels following allergen stimulation and CA treatment. * p<0.05 relative to BMMCs not treated with CA.

Taken together, these results suggest that a relatively unaffected MAPK signaling network coupled with a significant increase in Akt and NFκB activation may be the driving factors responsible for the observed increase in mediator gene expression and intracellular protein concentrations.

## Discussion

The current study provides a novel investigation into the anti-inflammatory mechanisms of the polyphenolic compound CA, in an IL-33 model of mast cell activation. Our results demonstrate for the first time that CA has the capacity to significantly reduce the secretion of key inflammatory mediators from IL-33 activated mast cells. Specifically, CA treatment was responsible for reducing the production of ROS, independent of any changes to the IL-33 specific inflammatory receptor-ome of mature mast cells. This impairment of early molecular events translated into a significant reduction in the release of key cytokines IL-6, IL-13, TNF, and chemokines CCL1, CCL2 and CCL3 (Fig. 2). Although secretion of these mediators was impeded, treatment with CA exacerbated IL-33 signaling through the Akt and NFκB pathway (Fig. 7), resulting in increased gene expression of *IL6*, *IL13* and *CCL3* (Fig. 3) in addition to intracellular protein levels of IL-6 and CCL3 (Fig. 4). Here, we also identify that IL-33 potentiation of allergen-induced mast cell activation is ameliorated following CA treatment, further supporting our previous investigation establishing CA as a potent anti-allergy compound (Fig. 3). Altogether, these results further characterize the versatility and therapeutic potential of CA treatment in various inflammatory pathologies, such as IL-33-mast cell associated diseases.

The use of CA as an anti-inflammatory compound has been investigated in several pathological contexts. In our previous investigation, we showed that CA effectively impaired IgE-mediated mast cell activation through a Syk-dependent mechanism^21^, resulting in a significant decrease in the release of pro-inflammatory mediators during both the early and late phases of allergic inflammation. Although this is the first study to investigate the effects of CA in a model of IL-33-mediated mast cell activation, IL-33 signaling shares highly conserved biochemical pathways with various TLR-family receptors, collectively resulting in the activation of the Myddosome signaling complex, which culminates in the production and release of a glut of pro-inflammatory mediators^24^. Similar to our current study which shows that CA inhibits the release of key cytokines such as IL-6 and TNF following IL-33 activation (Fig. 2), treatment with CA has been shown to reduce secretion of IL-6, TNF, IL-1β from Pam3CSK4 (TLR2 agonist)-stimulated bone marrow-derived macrophages ^25^ and TNF from LPS (TLR4 agonist)-stimulated RAW264.7 macrophages ^26^. Although the impairment of cytokine/chemokine release was expected as a result of our previous work and the current CA literature, we unexpectedly observed a significant increase in the gene expression levels of *IL6*, *IL13* and *CCL3* (Fig. 3), in addition to increased signaling through Myd88 (Fig. 6B), Akt and NFκB signaling nodes (Fig. 7). This not only differs from our previous work in another context which showed that 50 μM CA significantly reduced gene expression of *IL6, IL13,* and *CCL3* following IgE activation, but it also conflicts with the previously mentioned studies investigating the effects of CA on TLR signaling, where it was consistently observed that gene expression levels of key mediators such as IL-6 were significantly reduced.

The difference in inflammatory gene expression in our current study seems to stem from critical differences in the effects of CA on central signaling nodes. We previously observed in our mast cell model that treatment with CA abrogated downstream signaling through the Akt and NFκB pathways *via* a Syk-dependent mechanism^21^. Although it has been previously reported that IL-33 stimulation leads to the downstream phosphorylation and subsequent activation of Syk^8, 27^ , we were unable to observe any phosphorylation of Syk in our mast cell model following IL-33 and SCF activation (data not shown). In the absence of Syk as a central target, treatment with CA exacerbates mast cell signaling and the associated inflammatory response following IL-33 stimulation. CA has been well documented in various contexts of cell activation as an inhibitor of Myd88 accumulation^28^, downstream MAPK nodes ERK, JNK and p38^25,26,28^, in addition to NFκB proteins^25,26,28^ following TLR2, TLR4 activation and IgE-mediated mast cell activation^21^. Collectively, this increased signaling and subsequent gene expression in our IL-33 activated mast cells was responsible for the accumulation of IL-6 and CCL3 found within the cell that failed to be secreted (Fig. 5A, F). This intriguing data is suggestive that CA may target specific mechanisms responsible for the regulated secretion of pro-inflammatory mediators in activated mast cells, although further experiments are required to confirm this notion.

Protein trafficking mechanisms within a cell are comprised of a complex network of biochemical systems that are responsible for the regulated release of mediators upon various types of stimulation. Mast cells participate in several different forms of exocytosis that are dependant not only on the type of stimuli, but the timing of mediator release that follows cellular activation. These highly characterized secretory pathways, which include compound, multivesicular, piecemeal, and constitutive exocytosis^29,30^, are mainly involved in the early phase degranulation response (with the exception of the latter), but share some conserved features with the more poorly understood secretory pathways responsible for cytokine and chemokine release. These highly conserved features of exocytosis include members of Rab and Rho proteins, cytoskeletal proteins such as actin and tubulin, in addition to the cognate binding of vesicle and membrane bound SNARE proteins^31,32^. Rab proteins, the largest arm of the Ras-GTPase superfamily of proteins, have been implicated as central regulators of intracellular vesicle transportation, allowing for packaged proteins to be translocated to the cell membrane from the endoplasmic reticulum and golgi apparatus, to be released upon cellular activation^33^. An interesting feature of these proteins is they contain various sets of redox-sensitive motifs, providing an activation site for ROS to increase the exchange rate of GDP-to-GTP, ultimately promoting their activation^34^. Here, along with our previous report in an IgE-model of mast cell activation, we provide overwhelming evidence of the antioxidant capacity of CA, which significantly dampens stimuli-mediated ROS production (Fig. 1D-E) in various contexts. This overall reduction in ROS production within the cell could in turn reduce the activity of several Rab proteins and result in the stagnation of cytokine and chemokine movement, resulting in the accumulation within the cell, due to the lack of functional shuttling services (Fig. 5).

Although widely implicated in the early phase degranulation response, the importance of SNARE proteins in cytokine and chemokine release is becoming increasingly appreciated ^35^. The interaction of cognate SNARE proteins helps to facilitate the docking and subsequent fusion of secretory vesicles with the plasma membrane, allowing the release of vesicle-associated mediators ^36^. To test the capacity of polyphenols potentially impairing secretory mechanisms, an investigation by Yang et al^37^ looked at the effects of several polyphenol compounds on the formation or fusion of SNARE protein complexes. Despite the fact that the researchers focused on the early phase of mast cell/basophil activation, it was discovered that treatment with various polyphenols led to a reduction in the complex formation of syntaxin 4/SNAP-23/VAMP2 as well as the formation of syntaxin 4/SNAP-23/VAMP8^37^ complexes. The requirement of these SNARE protein complexes have been implicated in the release of several cytokines and chemokines associated with IgE and/or IL-33 activation, which includes the release of IL-6, TNF ^38^ and chemokines such as CCL5 ^30,31^. Although it has not currently been established that CA can disrupt the formation of these key SNARE complexes, this potential mechanism of action could explain why we saw an overall decrease in pro-inflammatory mediator release, despite an increase in specific genes and corresponding protein levels within the cell. This study warrants further investigation into the potential effects of CA on the various secretory mechanisms mast cells use, not only during the rapid expulsion of pre-formed mediators during degranulation, but also the regulated release of *de novo* synthesized cytokines and chemokines during allergen and IL-33-mediated mast cell activation.

Collectively, the data presented in this study further reveals the therapeutic potential of CA as a mast cell regulator in various contexts of inflammation. Following IL-33 activation, treatment with CA had a profound effect on reducing the release of pro-inflammatory cytokines and chemokines in the presence or absence of IgE-mediated co-stimulation, despite an increase in cell signaling and pro-inflammatory gene expression. These confounding results provide substantial evidence to further explore the underlying biological activity of CA in order to uncover the possible effects of this polyphenol within the highly intricate secretory mechanisms associated with mast cell activation. Regardless of the current unknowns, due to the ever-increasing evidence placing IL-33 as one of the main contributors of disease severity (allergy, asthma), this study not only provides additional evidence supporting the anti-allergy potential of CA, but also implicates the use of this polyphenolic compound as an anti-inflammatory approach to help combat the severity of IL-33-associated disorders.

## Authorship Contributions

RWEC and AJM conceived and designed the study. RWEC was assisted by JTM in carrying out experiments. RWEC analyzed the data. All authors contributed to writing, discussion and editing of the manuscript.

## Abbreviations

BMMC: bone marrow-derived mast cell
BSA: bovine serum albumin
CA: carnosic acid
CCL: CC chemokine ligand
cDNA: complimentary DNA
DCFH-DA, 2’ 7’: dichlorofluorescein diacetate
DMSO: dimethysulfoxide
DNP: dinitrophenyl
ELISA: enzyme-linked immunosorbent assay
ERK: extracellular signal-regulated kinase
FcR: Fc receptors
FITC: fluorescein isothiocyanate
HBSS: Hanks’ balanced salt solution
JNK: c-Jun N-terminal kinase
MAPK: mitogen activated protein kinase
MFI: mean fluorescence intensity
NFκB: nuclear factor kappa B
PBS: phosphate-buffered saline
PE: phycoerythrin
Pen/Strep: penicillin/streptomycin
p-NAG: p-nitrophenyl-N-acetly-β-D-glucosaminide PVDF, polyvinylidene difluoride
qPCR: quantitative polymerase chain reaction ROS, reactive oxygen species
SCF: stem cell factor
SNARE: Soluble N-ethylmaleimide-sensitive factor attachment protein receptors

